# G1 length dictates H3K27me3 landscapes

**DOI:** 10.1101/2023.12.05.570186

**Authors:** Abby Trouth, Kamesh Ravichandran, Philip R Gafken, Sara Martire, Gabriel E. Boyle, Giovana M. B. Veronezi, Van La, Stephanie J. Namciu, Laura A. Banaszynski, Jay F. Sarthy, Srinivas Ramachandran

## Abstract

Stem cells have lower facultative heterochromatin as defined by trimethylation of histone H3 lysine 27 (H3K27me3) compared to differentiated cells. However, the mechanisms underlying these differential H3K27me3 levels remain elusive. Because H3K27me3 levels are diluted two-fold in every round of replication and then restored through the rest of the cell cycle, we reasoned that the cell cycle length could be a key regulator of total H3K27me3 levels. Here, we propose that a fast cell cycle restricts H3K27me3 levels in stem cells. To test this model, we determined changes to H3K27me3 levels in mESCs globally and at specific loci upon G1 phase lengthening – accomplished by thymidine block or growth in the absence of serum (with the “2i medium”). H3K27me3 levels in mESC increase with G1 arrest when grown in serum and in 2i medium. Additionally, we observed via CUT&RUN and ChIP-seq that regions that gain H3K27me3 in G1 arrest and 2i media overlap, supporting our model of cell cycle length as a critical regulator of the stem cell epigenome and cellular identity. Furthermore, we demonstrate the inverse effect – that G1 shortening in differentiated cells results in a loss of H3K27me3 levels. Finally, in tumor cells with extreme H3K27me3 loss, lengthening of the G1 phase leads to H3K27me3 recovery despite the presence of the dominant negative, sub-stoichiometric H3.1K27M mutation. Our results indicate that G1 length is an essential determinant of H3K27me3 landscapes across diverse cell types.

## INTRODUCTION

Pluripotent stem cells must balance differentiation potential with their cellular identity [1]. Maintenance of cellular identity across multiple divisions is accomplished in part by the inheritance of parental chromatin states [2]. These states include the trimethylation of histone H3 lysine K27 (H3K27me3) deposited by Polycomb Repressive Complex 2 (PRC2). PRC2 deposits H3K27me3 across large chromatin domains that contribute to cell-type specific silencing [3]. The global distribution of H3K27me3 is cell-type specific [4]. Pluripotent stem cells have unique features in terms of H3K27me3 [5]: it is present at lineage specifying genes at low levels, leading to speculation that maintaining low levels of silencing might be enough to maintain stem cell state, at the same time providing stem cells with a lower barrier for differentiation. Stem cells must remain on the cusp of differentiation, requiring limited and reversible silencing. The mechanisms stem cells utilize to maintain low levels of H3K27me3 remain unclear, though a potential candidate arises when comparing how different cell types proceed through the cell cycle.

The cell cycle – consisting of the G1, S, and G2 phases followed by mitosis – is traversed to produce two identical daughter cells with the same chromatin landscape as the parent cell from which they arose. However, the DNA replication within the S phase challenges the levels of pre-existing epigenetic modifications like H3K27me3, as modified parental histones are divided amongst the two daughter strands in addition to the deposition of new, unmodified histones [6]. This dilution to approximately half the original H3K27me3 levels must be replenished before the subsequent replication to avoid a loss of silencing across multiple divisions. While every cell must overcome this dilution obstacle, cell cycle dynamics are highly heterogeneous across cell types, with notable differences between stem and differentiated cells. Stem cells speed through their cell cycle, with many examples where they forgo checkpoints or whole cycle phases. In the early embryonic stages of *Drosophila*, the cycle is 8 minutes long and composed of only the S phase and mitosis [7]. Serum-grown mouse embryonic stem cells (mESCs) divide in ∼11 hours [8], lacking a G1 checkpoint, shortening the G1 length [9]. mESCs have low levels of H3K27me3 compared to differentiated cells [10]. Thus, the time spent in the cell cycle could impact the global levels of H3K27me3 and heterochromatin in general.

We propose that the faster cell cycles observed in embryonic stem cells restrict H3K27me3 levels and, in turn, allow cells to differentiate more readily. To test this model, we determined changes to H3K27me3 levels in mESCs both globally and at specific loci upon G1 phase lengthening – accomplished by thymidine block or growth in the absence of serum (in the presence of two kinase inhibitors, PD0325901 and CHIR99021, the “2i medium” [11]). H3K27me3 levels in mESC increase with G1 arrest when grown in serum and are higher in 2i medium. Additionally, we observed via CUT&RUN and ChIP-seq that regions that gain H3K27me3 in G1 arrest and 2i media overlap, supporting our model of cell cycle length as a critical regulator in the stem cell epigenome and cellular identity. Furthermore, we demonstrate the inverse effect – that G1 shortening in differentiated cells results in a loss of H3K27me3 global levels via domain shrinking. Finally, in tumor cells with extreme H3K27me3 loss, lengthening of the G1 phase leads to H3K27me3 recovery despite the presence of the dominant negative, sub-stoichiometric H3.1K27M mutation. Our results indicate that G1 length is an essential determinant of facultative heterochromatin landscapes across diverse cell types.

## RESULTS

### Lengthening G1 increased global levels of H3K27me3 in mESC

We hypothesized that the low level of H3K27me3 present in serum-grown mESCs is due to the short G1 phase in these cells. To test this hypothesis, we first blocked mESCs in the G1/S phase with thymidine for 24 hours and then released the cells. We profiled H3K27me3 and H3 by immunoblots in asynchronous cells 24 hours after block (“0 hours”) and at time points up to 48 hours post-release. After a 24-hour block in G1/S, we observed a striking increase in H3K27me3 relative to H3 (**Figure 1A**). Notably, even up to 8 hours post-release, the H3K27me3 to H3 ratio remained significantly higher than asynchronous cells. Because unperturbed mESCs replicate in ∼11 hours [8], the persistent increase in H3K27me3 levels until 8 hours post-release indicates that one round of replication is insufficient to dilute H3K27me3 to pre-block levels. Therefore, lengthening the G1/S phase introduces a transient increase in H3K27me3 levels in mESCs. We next asked if the gain in H3K27me3 was proportional to the length of the G1/S phase block. We blocked mESCs at the G1/S phase with thymidine for 12 and 24 hours. The H3K27me3 levels increased proportionally with the length of the thymidine block, consistent with a model where the longer a cell spends in G1, the more H3K27me3 accumulated (**Figure 1B**).

**Figure 1.**
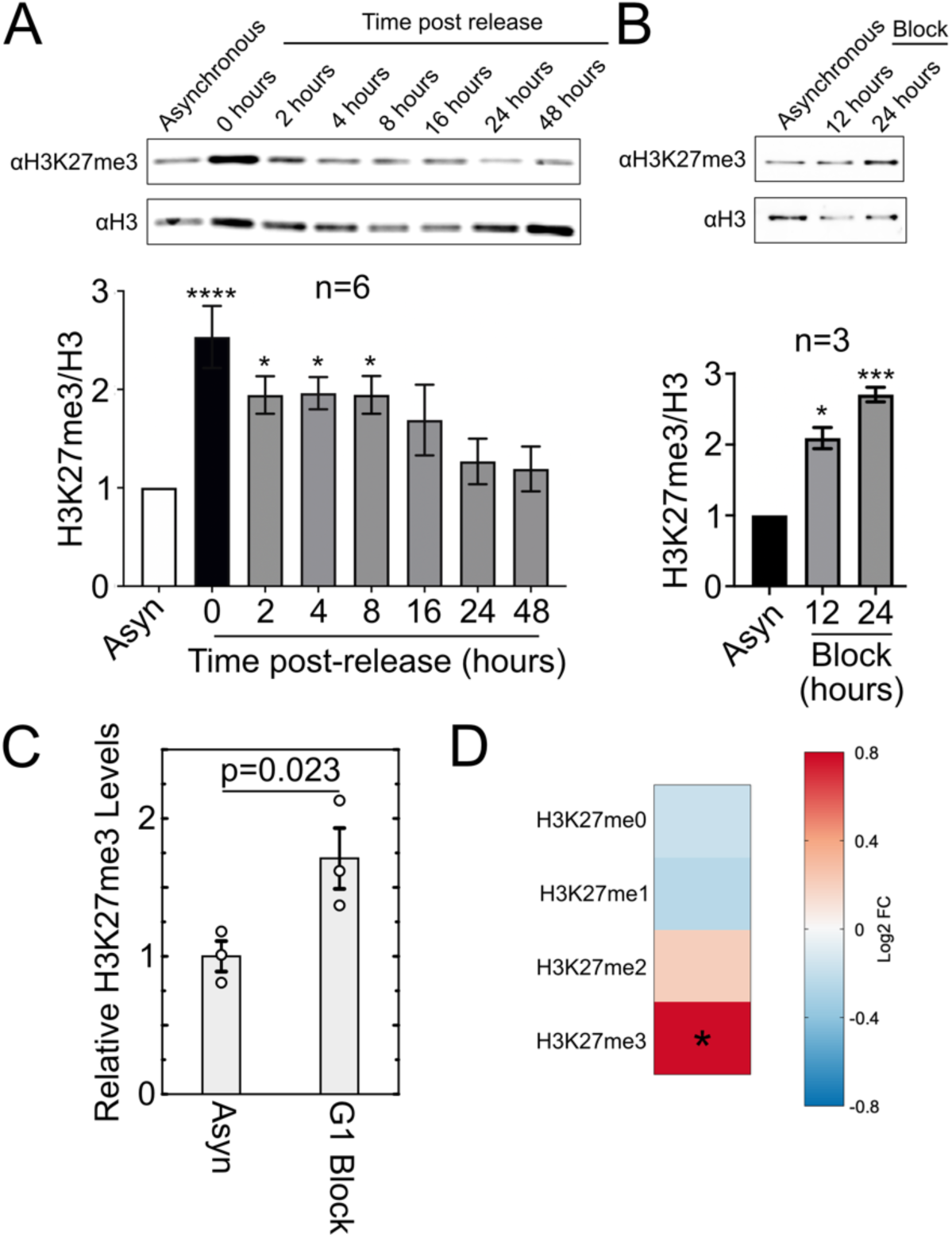
Global H3K27me3 levels in serum/LIF-grown mESCs increase with G1/S arrest. **A**) Immunoblot (top) and quantification (bottom) of the H3K27me3 modification relative to total H3 after G1/S arrest. Histones were acid extracted from mESCs treated with thymidine for 24 hours before release. Relative H3K27me3 levels are normalized to that of asynchronous cells. **B**) Immunoblot (top) and quantification (bottom) for H3K27me3 and total H3 on acid-extracted histones after 12 and 24 hours in G1/S arrest. Relative H3K27me3 levels are normalized to modification levels of asynchronous cells. **C**) Change in relative H3K27me3 levels determined via mass spectrometry across three replicates for cells that underwent double thymidine block with the second block lasting 20 hours. **D**) Heat map of average fold change across three replicates in H3K27 methylation states in G1 arrested cells vs. asynchronous cells. **E**) Heat map of fold change in methylation states for H3K36 and H3K9 after a 20-hour G1/S arrest for average values across the three replicates shown in (**C**). Values for each methylation state were normalized to values obtained from asynchronous cells. Asterisk (*) indicates p<0.05, p-values calculated using Student’s t-test with a two-tailed distribution.

To independently verify that H3K27me3 levels increase with the G1 lengthening, we analyzed histones by mass spectrometry after a double thymidine block to induce a G1 arrest. We performed a double thymidine block, where the second block was for 20 hours. We then purified histones by acid extraction and analyzed them by mass spectrometry (MS) to reveal all states of H3K27 methylation and quantify the change in H3K27me3 due to lengthened G1. Similar to immunoblots, we observed increased H3K27me3 in arrested cells (**Figure 1C**). We observed a decrease in H3K27me0/1 and an increase in H3K27me2/3 (**Figure 1D**). Thus, G1 lengthening results in a progression of methylation, converting unmethylated H3K27 and H3K27me1 towards H3K27me2 and H3K27me3. In summary, H3K27 methylation states correlated with G1 length in mESCs, with H3K27me3 showing the most substantial increase upon lengthening G1. This increase suggests that global epigenomic states in mESCs are influenced to a large extent by cell cycle length.

### G1 extension results in domain spreading and *de novo* domain formation

The global increase in H3K27me3 upon extension of G1 could be due to three scenarios – i) an increase in enrichment in existing H3K27me3 domains, ii) spread beyond the boundaries of existing domains to form larger ones, or iii) creation of new domains. To ask how the genome-wide distribution of H3K27me3 changed upon G1 extension, we performed CUT&RUN on H3K27me3 in triplicate after a single thymidine block for 8, 12, 16, and 20 hours, with asynchronous cells as control. We first defined domains for each time point, then compared domain definitions across time points to identify unique, non-overlapping segments that would make up the superset of domains across time points (**Figure 2A**). Among the six clusters obtained by k-means clustering, clusters 5 and 6 featured a gain in H3K27me3 and accounted for most segments of H3K27me3 that changed upon the thymidine block (41% bp of segments, covering a total of ∼87 million bp). Interestingly, clusters 1 and 2 exhibited a loss in H3K27me3, showing that, even though H3K27me3 goes up globally, some segments in the genome still lose H3K27me3. Thus, a more extended G1 phase rewires the genome-wide H3K27me3 landscape in mESC (**Figure 2B**).

**Figure 2.**
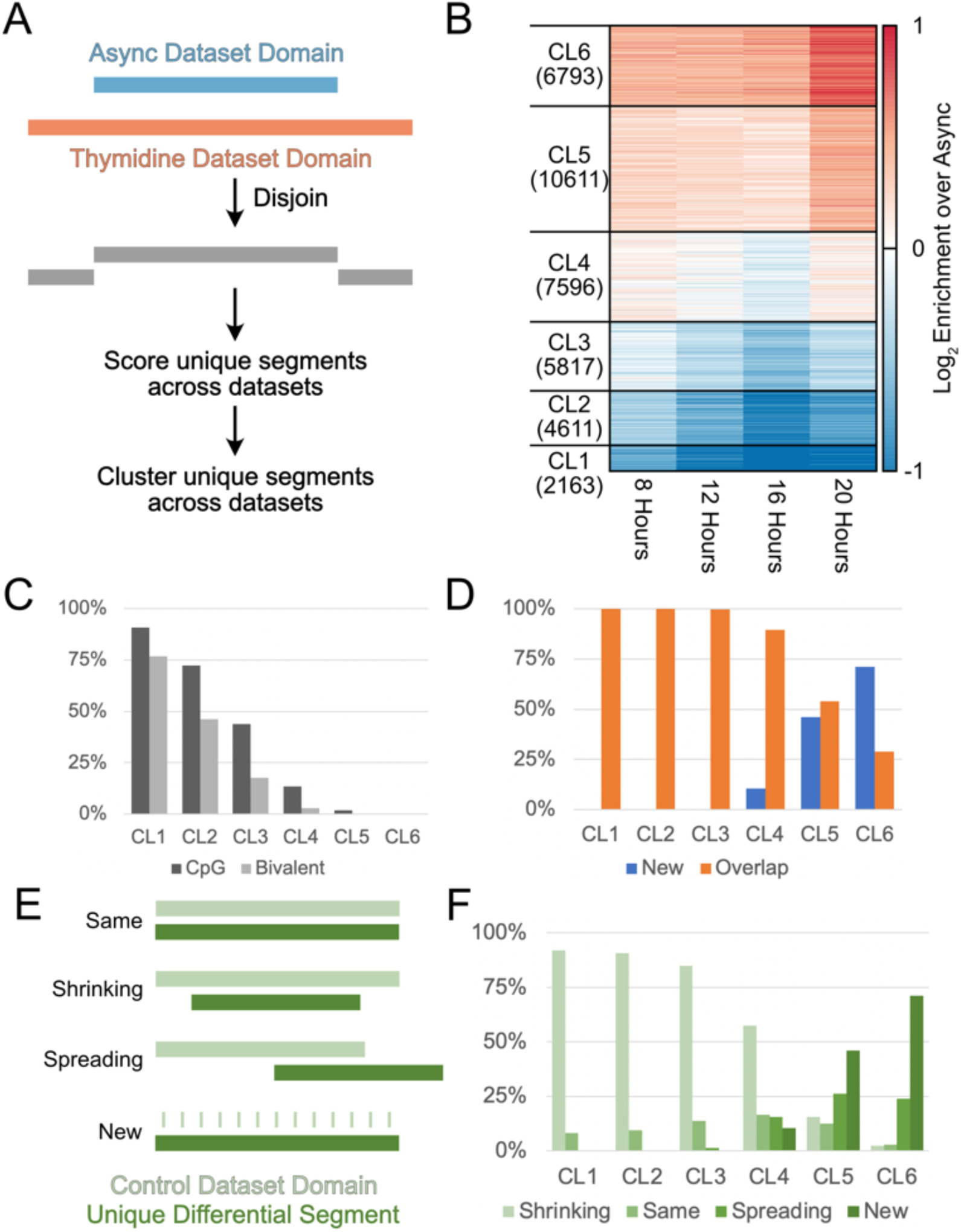
Locus-specific changes in H3K27me3 upon G1/S arrest. **A**) Schematic detailing how non-overlapping segments are defined by comparing H3K27me3 domains for asynchronous and thymidine-treated mESCs. **B**) Heatmap of H3K27me3 CUT&RUN enrichment at four time points of G1/S arrest after performing k-mean clustering with 6 clusters. Each horizontal line of the heatmap represents a unique segment determined by Disjoin. **C**) Percentage of segments in each of the 6 clusters containing CpG islands or bivalent promoters. **D**) Percentage of each of the 6 clusters that overlaps with H3K27me3 domains present in asynchronous mESCs. **E**) Schematic detailing four domain changes observed after thymidine arrest. Unique segments from thymidine datasets were classified as “same” when maintaining the same boundaries as an asynchronous domain, “shrinking” when one or more boundaries retract, “spreading” when one or more boundaries extend past those of the asynchronous domain, and “new” for domains not present in asynchronous cells. **F**) Percentage of each of the four domain changes in the 6 clusters.

PRC2 is known to act on two prominent, overlapping genomic features in mESCs [12]: CpG islands with low DNA methylation and bivalent promoters featuring both H3K27me3 and H3K4me3. Having clustered H3K27me3 domain changes in mESCs upon G1/S arrest, we next investigated the extent to which clusters overlapped with the two known markers of PRC2 activity. We observed a striking trend where the fraction of segments containing CpG islands/bivalent promoters monotonically decreased from cluster 1 to cluster 6 (**Figure 2C**). Notably, the two clusters with the most significant gain in H3K27me3-CL5/6 - show almost no overlap with CpG islands or bivalent promoters.

The lack of overlap between CpG islands/bivalent promoters and clusters 5 and 6 leads to two possibilities: i) the observed H3K27me3 increase in longer G1 phases may result from PRC2 spreading away from CpG islands and bivalent promoters or ii) the extended time in G1 allows for *de novo* nucleation at new sites that are not in proximity to these elements. To determine which mechanism is at play, we first asked how many segments in each cluster overlapped with H3K27me3 domains in asynchronous cells. In clusters 5 and 6, we observed new segments that did not intersect with domains in asynchronous cells, indicating *de novo* nucleation due to G1 extension (**Figure 2D**). We next categorized the segments that do overlap with domains in asynchronous cells as “same”/”shrinking”/”spreading” (**Figure 2E**). Clusters 1, 2, and 3 mainly feature shrinking domains due to thymidine treatment, whereas clusters 5 and 6 comprise some spreading and mostly new domains (**Figure 2F**). In summary, G1 extension results in more regions in the genome gaining H3K27me3 than losing it. The observed gain in H3K27me3 is partly facilitated by PRC2 spreading at previously defined domains. However, the modification is mainly gained by forming new domains that do not contain known PRC2 recruitment features. Our results suggest that extended G1 allows weaker nucleation sites to develop as domains, pointing to cell cycle time as an essential determinant in the genome-wide H3K27me3 landscape.

### Weak PRC2 nucleation sites drive H3K27me3 gain due to G1 extension

We next asked what the chromatin features at steady state were for regions that gain H3K27me3 due to G1 extension. We found the enrichment of four histone post-translational modifications to follow a trend from cluster 1 to cluster 6. H3K27me2 and H3K36me2 enrichment increased going from cluster 1 to cluster 6 (**Figure 3A, B**), whereas H3K27ac enrichment decreased from cluster 1 to cluster 6 (**Figure 3C**). Thus, clusters 5 and 6 could represent regions coated by H3K27me2 and H3K36me2 to silence cryptic enhancers [13–15]. H2AK119ub had a non-monotonic trend: it decreased from cluster 1 to cluster 4 but did not decrease further in clusters 5 and 6 (**Figure 3D, bottom**). At the center of the unique segments, the H2AK119ub enrichment goes up for cluster 6 after going down from clusters 1 to 5. Thus, clusters 5 and 6 feature significant levels of H2AK119ub, pointing to the possibility of segments in these clusters harboring weak nucleation sites for PRC1 and PRC2. We next asked if these four marks define different classes of chromatin in clusters 5 and 6 or if their trends correlate. We clustered the enrichment of these four marks using the k-means method (k=3) and observed a consistent trend across segments (**Figure 3E**). H3K27ac is uniformly low, and levels of H3K27me2, H3K36me2, and H2AK119ub correlate with each other. In summary, the sites that gain H3K27me3 due to extended G1 appear to be weak nucleation sites for PRC1 and PRC2, most probably driven by NSD1 [13]. We conclude they are weak nucleation sites due to an increasing trend for H3K27me2, opposite of the decreasing trend of H3K27me3 at steady state. Thus, PRC2 binds these regions and can modify H3K27 up to the dimethylation state but not efficiently to trimethylation due to short G1. When G1 is extended, there might be enough time for PRC2 to methylate these weak nucleation sites to the trimethylation level, leading to the observed significant increase in H3K27me3 with longer G1.

**Figure 3.**
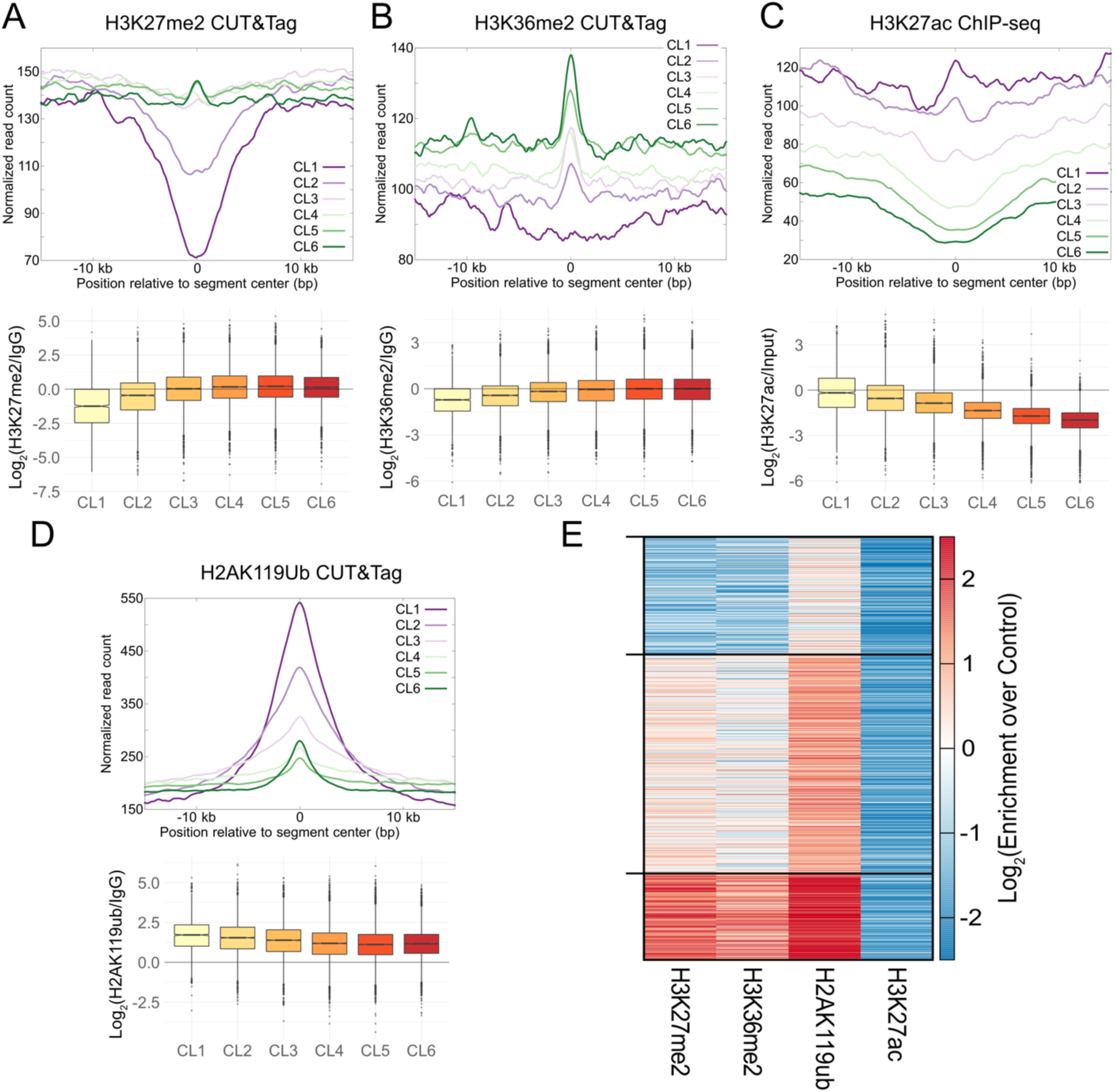
H3K27me3 gains due to longer G1 occur at weak nucleation sites. **A**) Enrichment of H3K27me2 CUT&Tag normalized read count averaged over unique segments in each thymidine cluster defined in Figure 2, plotted relative to centers of the unique segments (**top**). The log2 ratio of the H3K27me2 normalized read count at each segment to the IgG normalized read count is plotted as a boxplot for each thymidine cluster (**bottom**). **B**) Same as (**A**) for H3K36me2 CUT&Tag. **C**) Same as (**A**) for H3K27ac ChIP-seq. **D**) Same as (**A**) for H2AK119ub CUT&Tag. **E**) k-means clustering (k=3) of log2 enrichment of H3K27me2, H3K36me2, H2AK119ub, and H3K27ac for segments in thymidine clusters 5 and 6 that gain H3K27me3 plotted as a heatmap. H3K27ac is depleted in all clusters, and enrichment of H3K27me2, H3K36me2, and H2AK119ub correlate with each other.

### Regions that newly acquire H3K27me3 upon G1 extension overlap with 2i-specific H3K27me3 domains

Our experiments above were performed with mESCs grown in serum/LIF media. In these growth conditions, mESCs display a short G1 phase due to hyperphosphorylation of the Rb protein and subsequent lack of G1 checkpoints [9]. However, Rb is hypophosphorylated for mESCs grown in 2i media, and the G1 checkpoint is functional, resulting in a longer G1 [9]. Thus, our previous results suggest that 2i mESC should have higher global H3K27me3. We analyzed global H3K27me3 levels in mESCs from two sources, grown in 2i media or serum/LIF. We found higher levels of H3K27me3 in 2i mESCs compared to serum mESCs as described before [16] (**Figure 4A**).

**Figure 4.**
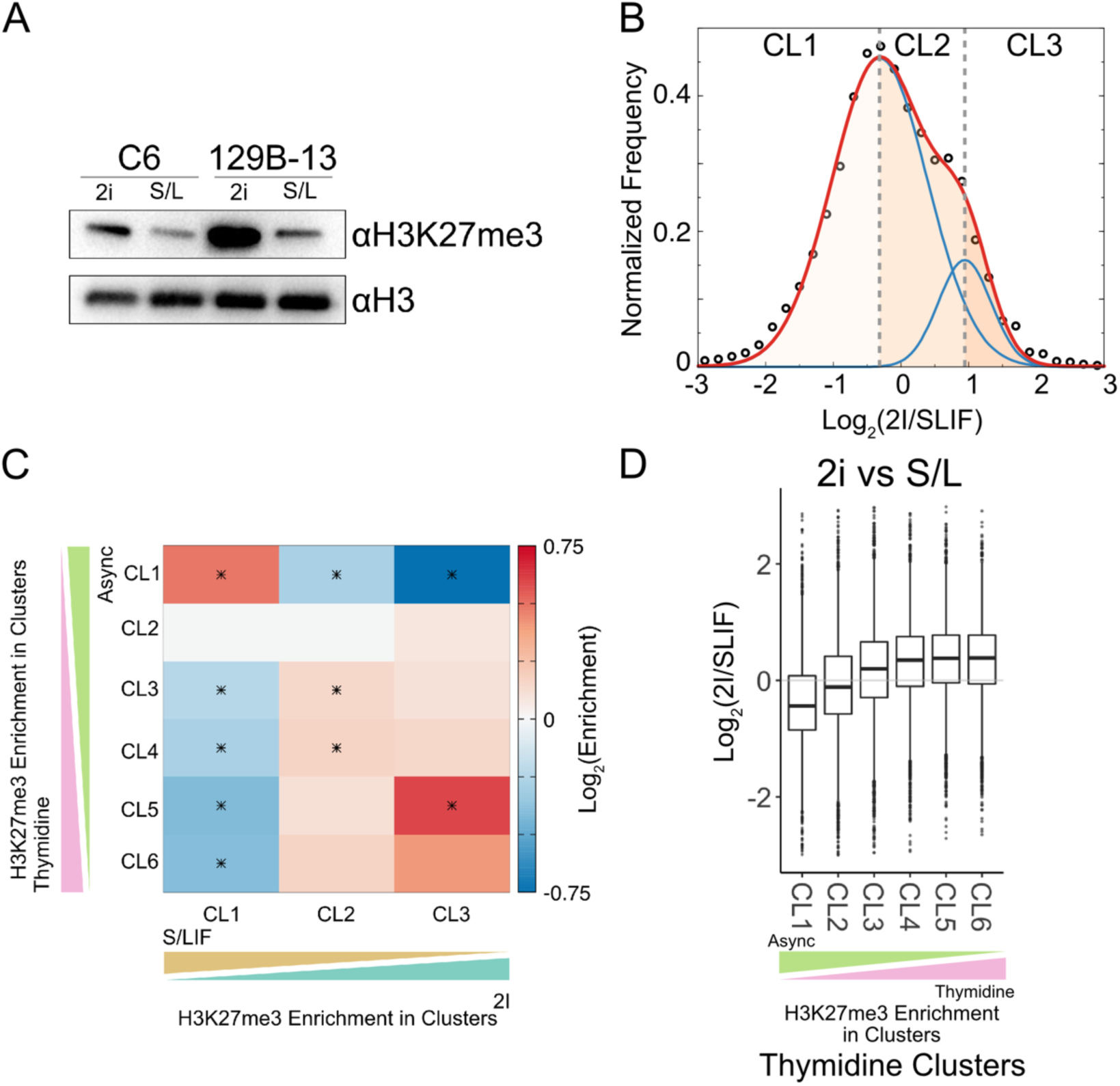
Regions with H3K27me3 gain in 2i-grown mESCs overlap with those in G1/S arrest of serum-grown mESC. **A**) Immunoblot for H3K27me3 modification of two mESC cell lines grown in either serum/LIF (S/L or SLIF) media or 2i media. **B**) Distribution of the log_2_ ratio of H3K27me3 levels in 2i versus serum cells for unique domain segments shown as black circles. The distribution was fitted with the sum of two normal distributions (plotted in red). The individual distributions are plotted in blue. The shaded regions correspond to the three groups based on the log2 ratio: more H3K27me3 in SLIF cells (CL1, leftmost region), equivalent H3K27me3 in SLIF and 2i cells (CL2, center region), and regions with more H3K27me3 in 2i cells (CL3, rightmost region). **C**) Extent of overlap of clusters of unique regions from G1/S block of serum grown cells and unique regions defined in (**B**). Asterisks denote statistical significance determined using hypergeometric test with multiple testing correction (corrected p<0.05) **D**) Log2 ratio of H3K27me3 enrichment of 2i mESC vs. serum mESC calculated at unique segments defined in Figure 2.

Next, we performed H3K27me3 ChIP-seq in both conditions to compare H3K27me3 changes at the domain level between 2i and serum/LIF mESCs. Similar to the analyses shown in Figure 2, we defined H3K27me3 domains genome-wide and then identified unique segments when comparing domains from 2i and serum/LIF cells. We observed a bimodal distribution in the ratio of H3K27me3 levels between these two conditions: the left-shifted distribution centered close to 0, indicating segments mostly similar between the two conditions; the right-shifted distribution centered close to 1, an average 2-fold increase in 2i compared to serum (**Figure 4B**). We defined 3 sets of regions – regions with higher H3K27me3 in serum/LIF (CL1), regions with equivalent H3K27me3 levels in both media (CL2), and regions with higher H3K27me3 levels in 2i (CL3) within the distribution (**Figure 4B**). If the length of the G1 phase was the underlying cause of the higher H3K27me3 observed in both 2i cells and thymidine-treated serum/LIF cells, then there should be significant overlap in regions that change in H3K27me3 levels between 2i cells and thymidine-treated serum/LIF cells. We overlapped clusters of segments in thymidine treatment with clusters from the 2i/serum/LIF comparison. We observe a significant enrichment between thymidine CL1 and 2i CL1, whereas a significant depletion between thymidine CL3-6 and 2i CL1. We also observe a significant enrichment between thymidine CL5 and 2i CL3 (**Figure 4C**). Thus, regions specific to serum/LIF cells compared to 2i cells overlap with regions that lose H3K27me3 upon G1 extension, and regions specific to 2i cells compared to serum/LIF gain H3K27me3 upon G1 extension. We can observe this also when we plot the Log2 ratio of H3K27me3 in 2i compared to SLIF at thymidine clusters; we observed an increase going from thymidine CL1 to CL6 (**Figure 4D**). In other words, we can produce an H3K27me3 landscape genome-wide similar to ground-state pluripotent stem cells (2i) simply by extending G1 using a thymidine block.

### Shortening G1 in differentiated cells reduces H3K27me3 levels

The global gain and locus-specific changes in H3K27me3 observed in serum/LIF-grown mESCs upon G1 phase extension illustrate the fundamental connection between the short cell cycle and low heterochromatin levels in stem cells. We next explored if this principle holds across cell types by asking if a differentiated cell would dilute H3K27me3 levels when forced to proceed through a short G1 phase. To shorten G1 in differentiated cells, we treated HEK293 cells with the Chk1 inhibitor Chiron-124 [17] for 48 hours – approximately the time for unperturbed cells to proceed through two divisions. After treatment, cells were stained with propidium iodide for cell cycle analysis by flow cytometry. We observed that most cells treated with DMSO were in G1 as expected, consistent with a long G1 phase (**Figure 5A**). In contrast and as expected, cells treated with Chiron-124 show a much more even distribution between all cell cycle phases as the inhibitor removes the G1 checkpoint (**Figure 5B**). We next determined the global H3K27me3 changes after Chiron-124 treatment. We purified histones by acid extraction and profiled H3K27me3 and H3 in HEK293 cells treated with DMSO or Chiron-124 for 48 hours by immunoblotting (**Figure 5C**). After 48 hours of Chiron-124 treatment, the cells possess approximately half of the global levels of H3K27me3 relative to H3 compared to the control DMSO-treated cells. These data support our model that G1 length significantly influences H3K27me3 levels.

**Figure 5.**
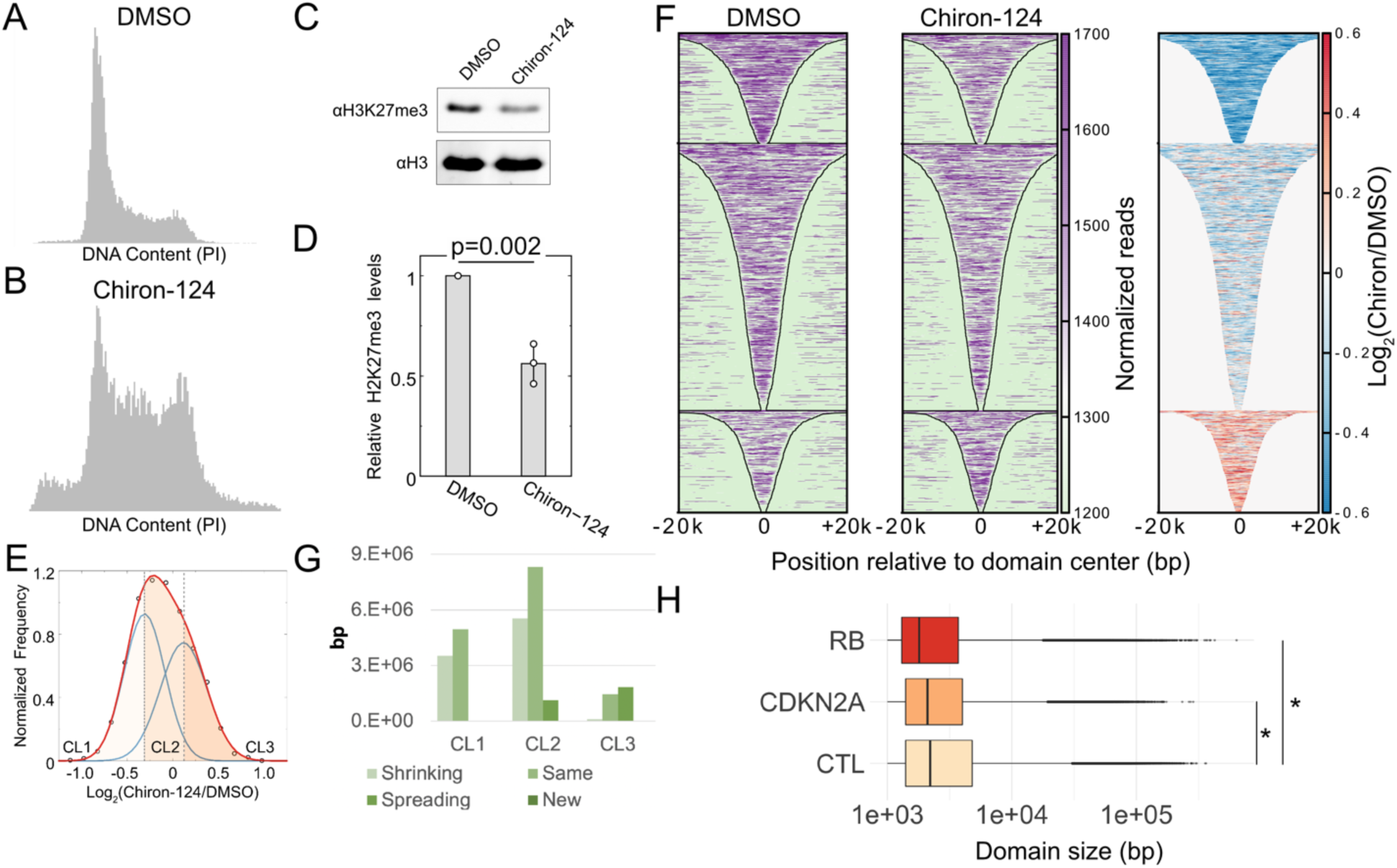
H3K27me3 domains in HEK293 cells change with accelerated cell cycle timing. **A**) Flow cytometry analysis of DNA content using propidium iodide fluorescence for HEK293 cells that were treated with DMSO for 48 hours. **B**) Same as (**A**) HEK293 cells treated with Chiron-124 for 48 hours. **C**) Immunoblot for H3K27me3 and H3 on acid-extracted histones after 48-hour treatment with Chiron-124. **D**) Quantification of the modification levels normalized to DMSO-treatment. **E**) Distribution of the log_2_ ratio of H3K27me3 levels in Chiron-124 treatment compared to the DMSO treatment for unique domain segments. Data points are shown as black outlined circles; the sum of the two normal distributions is plotted as red; individual normal distributions are plotted as blue lines. The shaded regions correspond to the three groups based on the log2 ratio: loss of H3K27me3 upon Chiron-124 treatment (CL1, leftmost region), equivalent H3K27me3 in Chiron-124 and DMSO treatment (CL2, center region), and regions with more H3K27me3 in Chiron-124 treatment (CL3, rightmost region). **F**) Heatmap of H3K27me3 enrichment at domains defined in DMSO treated cells for DMSO treatment (left) and Chiron-124 treatment (middle), and the log2 ratio of H3K27me3 within domains for Chiron-124 over DMSO (right). **G**) Percentage of each of the four domain changes (shrinking, same, spreading, and new) observed in the three clusters present in the bimodal distribution of H3K27me3 for Chiron-124 versus DMSO cells. **H**) Distribution of H3K27me3 domain sizes across ENCODE cell lines with inactivated Rb or CDKN2A deletion or neither. Domain sizes were determined from SEGWAY annotations from ENCODE. *p< 2.2e-16 by Wilcoxon rank sum test.

To profile locus-specific changes due to Chiron-124, we performed H3K27me3 CUT&Tag after 48 hours of treatment. We defined genome-wide H3K27me3 domains and generated unique segments from domain comparison between the treatment and control samples. To visualize H3K27me3 changes within these unique segments, we plotted the distribution of the log_2_ ratio of H3K27me3 levels at these unique segments in Chiron-124 versus DMSO-treated cells (**Figure 5E**). This distribution was left shifted relative to zero, indicating a global loss in H3K27me3, reflecting results obtained by immunoblot. We fit a bimodal distribution and defined three clusters based on their specific H3K27me3 change after G1 shortening – regions with lower H3K27me3 in Chiron-124 (CL1), regions with equivalent H3K27me3 between Chiron-124 and DMSO (CL2), and regions with higher H3K27me3 in Chiron-124 (CL3). Domains that intersected with CL1 segments featured a loss in H3K27me3 across their length upon Chiron-124 treatment and also appeared to shrink from their boundaries (**Figure 5F**). Next, we categorized segments within the three clusters based on their pattern of H3K27me3 change in short G1 phase cells compared to asynchronous (**Figure 5G**). CL1 and CL2, which cover much more base pairs than CL3, comprise segments that have shrunk compared to their corresponding domains in DMSO-treated cells. Thus, the global loss in H3K27me3 is manifested as the shrinking of individual domains on the genome, pointing to less time for PRC2 to spread the modification under a short G1 phase. In summary, in differentiated cells, shortening G1 results in loss of H3K27me3, showing that cell cycle length influencing global H3K27me3 levels might be applicable across cell types.

### Weakening of the G1 checkpoint correlates with smaller H3K27me3 domains

Inactivation of retinoblastoma protein (Rb) and cyclin-dependent kinase inhibitor 2A (CDKN2A) occurs frequently in cancer [18, 19]. Inactivation of these proteins is expected to weaken the G1 checkpoint, and we hypothesized that these perturbations would affect H3K27me3 domains due to reducing G1 length. To test this hypothesis, we defined H3K27me3 domains using SEGWAY [20] data from ENCODE [21] for cell lines that had inactivated Rb (HeLa, WERI-Rb-1) or had a CDKN2A deletion (A673, Panc1, SJSA1, and MCF-7), or neither (A549, HepG2, K562, SK-N-SH, Karpas-422, MM.1S, and PC-3). We coalesced segments defined as “FacultativeHet” by SEGWAY whose ends were within 500 bp of similar segments and then plotted the size distribution of H3K27me3 domains across the three categories. We observed the loss of CDKN2A and Rb to result in significantly smaller domains, with Rb loss having a bigger effect than CDKN2A (**Figure 5H**). Thus, complementary to the abrogation of the G1 checkpoint using Chiron-124 treatment, genetic weakening of the G1 checkpoint also results in smaller H3K27me3 domains, possibly due to shorter G1 in these cells.

### Recovery of H3K27me3 in DMG cells with the H3K27M mutation

Our results in mESCs and HEK293 cells point to the importance of the length of the G1 phase in maintaining low H3K27me3 levels genome-wide and ultimately preserving cellular identity. Our results also point to an unexpected therapeutic avenue in modulating heterochromatin in diseases where aberrant heterochromatin is associated with pathogenicity. One such disease is diffuse midline glioma (DMG), where children present with CNS tumors bearing an H3K27M mutation in one of the multiple copies of H3, resulting in a dominant negative effect of a global loss in H3K27me3 [22]. The substoichiometric H3K27M can lead to a spreading defect in PRC2 [23]. If H3K27me3 levels are substantially lowered in DMG cells due to reduced PRC2 activity, increasing cell cycle time could rescue H3K27me3 levels by giving PRC2 more time to modify H3 even at reduced efficiency. The CDK4/6 inhibitor Palbociclib restricts cells in the G1 phase [24]. We asked if treatment with Palbociclib, by extending G1, would result in increased levels of H3K27me3 in SU-DIPG-IV cells. H3K27me3 levels but not H3K27M levels increased with 72-hour Palbociclib treatment (**Figure 6A**). To investigate changes in H3K27me3 distribution after treatment with Palbociclib, we performed CUT&Tag in triplicate on H3K27me3 after 72-hour Palbociclib treatment. As previously described, domains were defined for control and Palbociclib and then compared with each other to identify unique segments. The distribution of the log_2_ ratio of H3K27me3 enrichment between Palbociclib and DMSO-treated cells showed a clear bimodal distribution that could be fitted with two normal distributions (**Figure 6B**). The mean of the left-shifted distribution was ∼0, whereas the mean of the right-shifted distribution was ∼0.7. This allowed us to separate the segments into three groups based on the means of the two underlying distributions: regions that have a minor loss in H3K27me3 (Group 1), regions that have a minor gain in H3K27me3 (Group 2), and regions that have a substantial gain H3K27me3 (Group 3) after Palbociclib treatment (**Figure 6B, D**). Representative regions with H3K27me3 gain from Group 3 show a notable increase within known Polycomb domains present in the SU-DIPG-IV cells in combination with H3K27me3 extending beyond the established domain borders (**Figure 6C**). Indeed, the majority of the unique segments within Group 3 correspond to spreading from domain boundaries defined in the control treatment (**Figure 6E**). Though the direct contribution of increased H3K27me3 spreading to cytostatic effects of Palbociclib in DMG cells is unknown, it is intriguing to note that extending G1 in these cells leads to increased H3K27me3 levels. As DMGs are defined by the co-occurrence of the H3K27M oncohistone and pathologically low levels of H3K27me3, cell cycle modulation may be a promising epigenetic therapy in this and other PRC2-deranged malignancies.

**Figure 6.**
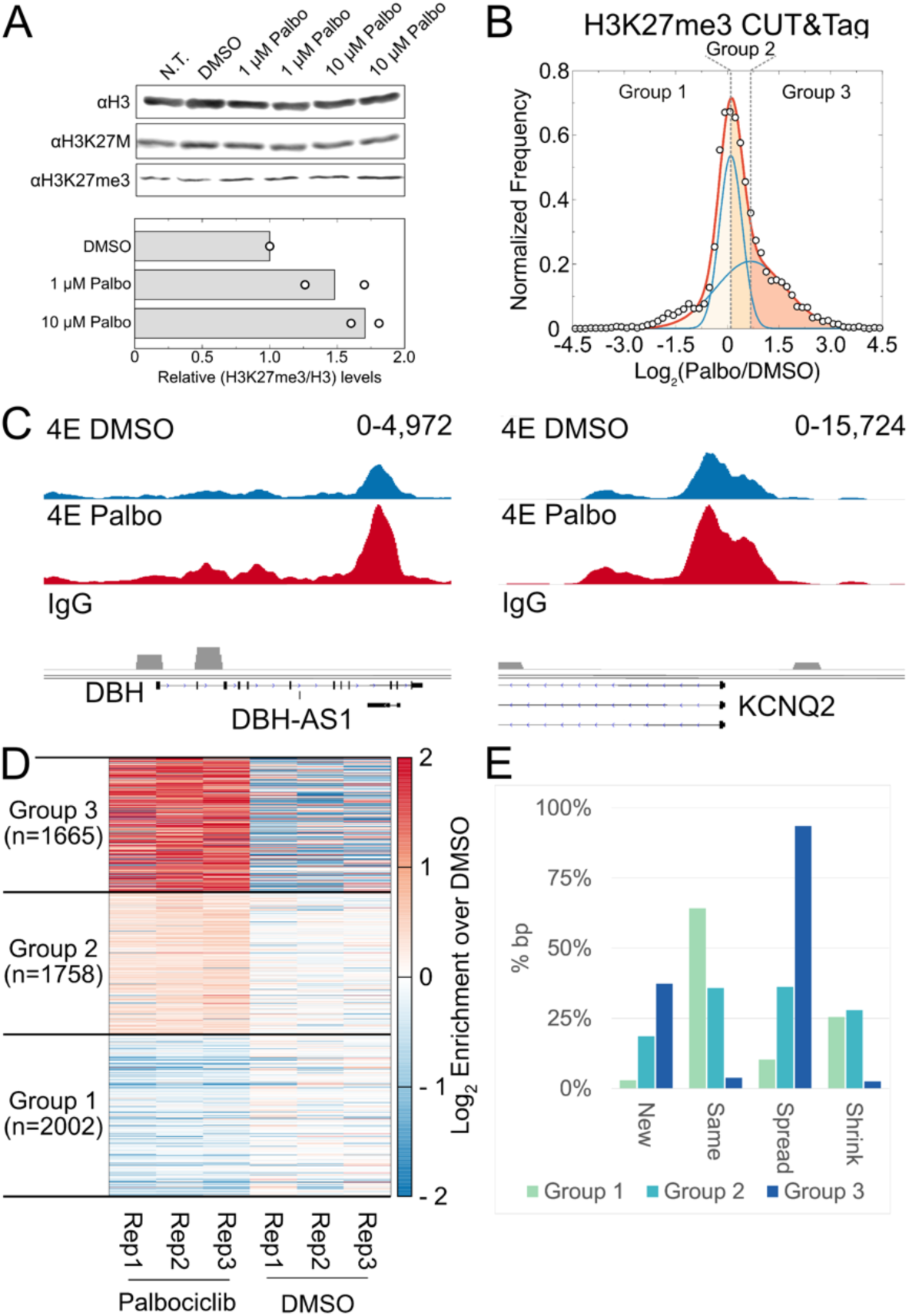
H3K27me3 domains are restored upon G1 arrest in patient-derived H3K27M mutant DMG cell lines. **A**) Immunoblot for H3, H3K27M, and H3K27me3 after 72-hour treatment with Palbociclib (top). Quantification of the modification levels normalized to DMSO-treatment (bottom). **B**) Plot of log_2_ ratio of H3K27me3 levels in Palbociclib compared to DMSO-treated SU-DIPG-IV cells for unique domain segments. The distribution was fitted with the sum of two normal distributions (plotted in red). The individual distributions are plotted in blue. The shaded regions correspond to the three groups based on the log2 ratio: loss of H3K27me3 upon treatment with Palbociclib (CL1, leftmost region), equivalent H3K27me3 in Palbociclib and DMSO treatment (CL2, center region), and regions with more H3K27me3 in cells treated with Palbociclib (CL3, rightmost region). **C**) Representative H3K27me3 domains from the H3K27me3 gain CL3. **D**) Heatmap of H3K27me3 CUT&Tag enrichment for SU-DIPG-IV cells treated with Palbociclib compared to DMSO vehicle control for the 3 clusters defined in (**A**). **E**) Percentage of each of the five domain changes observed in the 3 clusters defined in (**B**). Most domains present in CL1 after Palbociclib treatment maintain the same boundaries as observed in DMSO vehicle control samples. CL3 however, which presents the highest log_2_ enrichment of H3K27me3 after Palbociclib treatment, is mostly composed of domains with spreading boundaries that extend past those observed in DMSO control samples.

## DISCUSSION

Here, we have demonstrated the interplay between H3K27me3 and the cell cycle in human and mouse cells and shown that the length of the G1 phase influences the distribution and enrichment of H3K27me3 genome-wide. Upon manipulating the G1 phase, we observe domain-level changes of H3K27me3 spreading, shrinking, and *de novo* nucleation. Our results suggest that the cell cycle length plays a major role in shaping the epigenomic landscape, as observed in theoretical models [25].

In mESCs, G1/S arrest via thymidine block leads to global gain in H3K27me3 and domain-specific changes. These manipulations to the G1 phase were achieved in mESCs grown in serum, and later comparisons were made with mESCs grown in 2i medium. Serum-grown mESCs have hyperphosphorylated Rb and lack a G1 checkpoint, making their G1 short, whereas 2i mESCs have an intact G1 checkpoint, leading to a significantly longer G1 than serum mESCs [9]. Notably, the domains gained upon G1 extension in serum mESCs mirror those gained in 2i mESCs, underlining that the gain in H3K27me3 in 2i mESCs could be explained by their longer G1. The distribution of H3K27me3 in embryonic stem cells is crucial for both initial commitment and maintenance of cell identity, and changes in cell cycle length through development could provide a potential mechanism for rapidly changing the H3K27me3 landscape.

Rapid cell cycles in early development are a common theme in metazoans, making our observed influence of this cell cycle speed on the H3K27me3 landscape widely applicable. In *Drosophila* embryos, the establishment of the repressive chromatin modification H3K9me3 is influenced by changes to interphase duration that allow for cumulative recruitment of the methyltransferase Eggless [26]. Thus, constitutive and facultative heterochromatin may have a similar dependence on cell cycle timing for establishment during early development. While these results support our model of cell cycle timing as a regulator of global epigenomics in mESCs, there are still unanswered questions about how histone modifications other than H3K27me3 respond to G1 phase extension. Determining whether competition may exist between histone-modifying enzymes due to changes to the cell cycle would be an interesting future avenue of research.

We observed the enrichment of H3K27me3 at new domains that do not overlap with known PRC2 recruitment features, while there is a notable loss of H3K27me3 enrichment within CpG islands and bivalent domains. Sequence and protein interactions may aid PRC2 recruitment to the newer, weaker nucleation sites exposed by longer G1. Similarly, changes in local methyltransferase and demethylase activity could help restructure the epigenome as the stem cell changes its cell cycle in preparation for differentiation.

In support of G1 length as a major determinant of global H3K27me3 levels, forcing HEK293 cells to proceed faster through the G1 phase causes a loss of H3K27me3. Our results agree with the previous finding that most global H3K27me3 recovery occurs in G1 in differentiated cells [6]. An acceleration in the cell cycle also occurs during T-cell activation. Upon exposure to an antigen, CD8^+^ T cells undergo rapid proliferation controlled by the phosphorylation state of Rb and, therefore, the duration of the G1 phase [27]. However, it is unknown how the heterochromatin landscape responds to these sudden rapid cell cycles: the cells could either compensate by upregulating PRC2 function, or major changes in the heterochromatin landscape could accompany T-cell activation. The fact that activating mutations in Ezh2 leads to follicular and diffuse B-cell lymphoma at a stage where the B-cell cell cycle shortens points to the possibility that cell cycle shortening during the activation of immune cells could lead to heterochromatin remodeling [28].

Finally, we observed increased H3K27me3 levels in DMG cells with the dominant negative H3.1K27M mutation when treated with the selective CDK4/6 inhibitor Palbociclib. Nucleosomes containing mutant H3K27M can directly interact with PRC2 to prevent PRC2 spreading [29]. Our results support H3.1K27M cells displaying a PRC2 spreading defect that can be partially compensated by lengthened G1. Our results would also suggest a loss of H3K27me3 in cancers that lose the G1 checkpoint via mutations in Rb or deletions in Cdkn2a. In these tumors, optimal PRC2 function might be critical to maintaining H3K27me3 under fast cell cycles similar to serum mESCs, pointing to PRC2 as a potential target in these tumors. In summary, the cell cycle could control H3K27me3 dynamics in disparate systems, such as early development and tumor growth.

## AUTHOR CONTRIBUTIONS

S.R. conceptualized this study. K.R. performed immunoblots, sample preparation for mass spectrometry, and mESC differentiation. K.R. and A.T. performed flow cytometry and CUT&RUN in mESC. A.T. performed flow cytometry, immunoblots, and CUT&Tag in HEK293 cells. S.M. and L.A.B. performed immunoblots and ChIP-seq for 2i and serum-grown mESC. P.G. performed mass spectrometry. G.M.B.V. and V.L. performed CUT&Tag for H3K27me2, H3K36me2, and H2AK119ub. J.S. performed flow cytometry, immunoblots, and CUT&Tag in DMG cells. S.R. performed the data analysis. A.T. wrote the original draft of the manuscript, and S.R. edited it. All authors reviewed and approved the manuscript.

## ACKNOWLEDGEMENTS

This work was supported by the RNA Bioscience Initiative, the University of Colorado School of Medicine, and NIH grant R35GM133434 (S. R.), R35GM124958, R01HD109239, ACS 134230-RSG-20-043-01-DMC, and Welch Foundation I-2025 (L. A. B.). We thank Lisa Jones and Chenwei Lin for their assistance with mass spectrometry data collection and analysis. The Fred Hutch Proteomics & Metabolomics Shared Resource is supported by the Fred Hutch/University of Washington Cancer Consortium (P30 CA015704), and the M.J. Murdock Charitable Trust funded the Orbitrap Fusion instrument used in this research. SU-DIPG-IV cells were generously provided by the laboratory of M. Monje, Stanford University, and the laboratory of James Olson is acknowledged for assistance with cell culture. Shruti Bhise and Sami Kanaan are acknowledged for assistance with flow cytometry, and Ekaterina Babaeva for Western blotting. We thank Aaron Johnson and Olivia Rissland for critically reading the manuscript.

## METHODS

### Mammalian Cell Culture

All cell lines used in this work were tested for mycoplasma and confirmed to be negative for each experiment. Experiments utilized mouse embryonic stem cell line E3 (mESCs; Gates Biomanufacturing Facility at University of Colorado Denver), human embryonic kidney 293 cell line (HEK293; Fred Hutchinson), and patient-derived K27M SU-DIPG-IV (IV, H3.1K27M; J. Olson and J. Sarthy laboratories).

mESCs were cultured under feeder-free conditions on 0.1% gelatin-coated T-25 flasks in knockout DMEM (Life Technologies), supplemented with 20% embryonic stem cell-qualified fetal bovine serum (FBS; Life Technologies), 2 mM L-glutamine (Life Technologies), 0.1 mM non-essential amino acids (Life Technologies), 100UmL-1 penicillin/streptomycin (Life Technologies), 0.05 mM β-mercaptoethanol (Sigma), and 1000UmL-1 ESGRO mouse leukemia inhibitory factor medium supplement (LIF; Millipore).

For the comparison of serum/LIF and 2i conditions, Mouse embryonic stem cell lines (mESCs) were cultured on gelatin-coated plates at 37 C with 5% CO2 under either standard Serum/LIF conditions (KO-DMEM, 2 mM Glutamax, 15% ES grade fetal bovine serum, 0.1 mM 2-mercaptoethanol, 1x Pen/Strep, 1x NEAA and leukemia inhibitory factor (LIF)) or in 2i media (1:1 mix of DMEM/F12 and Neurobasal medium including N-2 supplement, B-27 supplement, 0.1 mM 2-mercaptoethanol, 2 mM glutamax, LIF, 3 µM CHIR99021 and 1 µM PD0325901).

HEK293 cells were cultured in DMEM (Life Technologies), supplemented with 10% FBS (Life Technologies) and 100 U/mL penicillin/streptomycin (Life Technologies). Cell lines were grown at 37°C with 5% CO_2_ in a humidified Forma Steri-Cycle CO_2_ incubator (ThermoFisher Scientific) and subcultured approximately every 2 days to maintain exponential growth. To harvest cells, a 0.25% trypsin-EDTA solution (Gibco) was applied for 3-4 minutes for mESCs and 1 minute for HEK293 cells. Detached cells were centrifuged at 2000 rpm and 4°C for 5 minutes and washed with 1x phosphate-buffered saline (PBS pH 7.4; Gibco) for use downstream.

The SU-DIPG-IV cell line had targeted sequencing to confirm H3 mutational status and was cultured as previously described [30].

### Thymidine Block for G1/S Arrest

For experiments utilizing thymidine to block cells in the G1/S phase, a final concentration of 2 mM thymidine (Sigma) was administered. The block-and-release experiments were performed by treating mESCs with thymidine for 24 hours, followed by a wash with 1x PBS and the addition of fresh medium to cells. Samples were taken for downstream use immediately after release and then again 2-, 4-, 8-, 16-, 24-, and 48-hours post-release. To determine how the length of the G1/S block affected global H3K27me3, mESCs were treated with thymidine for a total of 12 and 24 hours before being harvested for downstream use. The double-block experiments were performed by first treating mESCs with thymidine for 24 hours, releasing them for 7 hours via 1x PBS wash and recovery in a fresh medium, followed by a second block for 20 hours. In preparation for CUT&RUN, mESCs were treated with thymidine for 8, 12, 16, and 20 hours before being harvested together with asynchronous cells as control. These samples were then immediately processed for CUT&RUN.

### Chiron-124 Treatment for G1 Shortening

For experiments utilizing Chir-124 (MedChemExpress) to shorten the G1 phase for differentiated cells, a final concentration of 0.5 µM was administered to HEK293 cells. To determine how G1 shortening affected global H3K27me3, cells were treated with Chir-124 for 48 hours and then harvested for flow cytometry as described below. To determine domain-level H3K27me3 changes in HEK293 cells after G1 shortening, cells were treated with Chir-124 for 48 hours and then immediately processed for CUT&Tag.

### Palbociclib Treatment for G1 Arrest

For experiments utilizing Palbociclib (Cell Signaling Technologies) to arrest SU-DIPG-IV cells in the G1 phase, cells were treated for 72 hours before harvesting CUT&Tag experiments. For CUT&Tag profiling, cells were treated with 10 µM Palbociclib before being harvested for immediate use.

### Flow Cytometry for Cell Cycle Gating

Cells were harvested using a trypsin solution as detailed above and washed with 1 mL 1x PBS before being pelleted at 2000 rpm and 4°C for 5 minutes. The cells were then resuspended in 70% ethanol via gentle vortexing and incubated overnight at −20°C. Cells were pelleted at 2000 rpm and 4°C to remove ethanol for 5 minutes. Cells were resuspended in 500 µL Krishan stain [31] (9.25 mM sodium nitrate, 69 uM prodium iodide, 0.01% NP40, & 2 mg RNaseA) and incubated overnight at −20°C. Samples were analyzed using a Beckman Coulter FC500 flow cytometer. Cell gating was performed using the FlowJo software (BD Life Sciences).

### Histone Purification via Acid Extraction

Histones were purified from flash-frozen cells using a standard acid extraction protocol adapted from Shechter *et al.* [32]. Cells were thawed on ice and then lysed in hypotonic lysis buffer (10 mM Tris-HCl pH 8.0, 1 mM KCl, 1.5 mM MgCl_2_, and 1 mM DTT). Intact nuclei after lysis were pelleted by centrifugation at 10,000 x g and 4°C for 10 minutes before being resuspended in 0.4N H_2_SO_4_ and incubated at 4°C overnight. After incubation, nuclear debris after lysis was pelleted via centrifugation at 16,000 x g and 4°C for 10 minutes. Histones were precipitated via the dropwise addition of 150 µL 100% trichloroacetic acid and then incubated on ice for 30 minutes. The precipitated histones were pelleted via centrifugation at 16,000 x g and 4°C for 10 minutes, washed once with cold acetone, and then air-dried at room temperature for 20 minutes. The final histone pellet was then dissolved in 100 µL DI water. The concentration of histones in each sample was determined using the BCA protein assay kit (ThermoFisher Scientific).

### Immunoblotting Analysis

Acid-extracted histones were prepared in 1 µg samples, separated by electrophoresis using a 15% SDS-polyacrylamide gel, and transferred to a nitrocellulose membrane at room temperature. Whole-cell extracts were separated using a 4-20% Tris-Glycine polyacrylamide gel (Invitrogen) and transferred to a nitrocellulose membrane. Histones isolated from mESCs and HEK293 cells treated with thymidine and Chir-124, respectively, were probed with 1:1000 dilutions of antibodies against H3K27me3 (#9733, Cell Signaling Technologies) and total H3 (#18521, Abcam). Following overnight primary antibody incubation, membranes were probed with 1:10000 dilutions of IRDye 800CW Goat anti-Mouse IgG (926-65010, Fisher Scientific) and IRDye 800CW Goat anti-Rabbit IgG (926-32211, Fisher Scientific) and imaged with a Sapphire Biomolecular Imager (Azure Biosystems) using 784/BP22 and 658/BP22 membrane focus settings. Signals for each blot were quantified using the ImageJ software [33].

### Mass Spectrometry Analysis of H3K27me3

Aliquots corresponding to 10 µg of enriched histones in water were placed in Eppendorf tubes and taken to dryness by vacuum centrifugation. Histone derivatization and digestion were carried out similarly to that described by Sidoli *et al.* [34]. Dried samples were resuspended in 20 µL of 100 mM ammonium bicarbonate, followed by the sequential addition of 5 µL acetonitrile, 5 µL propionic anhydride, and 14 µL ammonium hydroxide. The derivatization reaction was incubated at 37°C with mixing for 15 minutes and then dried by vacuum centrifugation. The derivatization reaction was repeated once and dried by vacuum centrifugation to approximately 10 µL. Enzymatic digestion was carried out by adding 50 µL of 1 M ammonium bicarbonate to neutralize the solution and 1 ug of trypsin, after which the digestion reaction was incubated overnight at 37°C with mixing. Samples were dried to approximately 20 µL, and peptide derivatization was carried out twice, as previously detailed, ending with dried samples. The samples were resuspended in 20 µL of 0.1% trifluoroacetic acid, desalted with C18-micro ZipTips (Millipore), and taken to dryness. LC-ESI-MS/MS (liquid chromatography coupled to electrospray ionization tandem mass spectrometry) was carried out with an Orbitrap Fusion (ThermoFisher Scientific) mass spectrometer using an instrument configuration and data-independent acquisition (DIA) for bottom-up proteomics as described by Karch *et al.* [35]. DIA data were analyzed using EpiProfile v2.0.

### CUT&RUN

Genome-wide H3K27me3 domain changes were determined using the CUT&RUN protocol adapted from Hainer & Fazzio [36]. Cells were harvested via 0.25% trypsin-EDTA solution and counted using 0.4% trypan blue stain and Countess automated cell counter (Invitrogen). A single-cell suspension was then prepared to achieve 0.5 x 10^6^ cells in 1 mL cold nuclear extraction buffer (20 mM HEPES-KOH pH 7.9, 10 mM KCl, 0.5 mM spermidine, 0.1% Triton x-100, & 20% glycerol). Isolated nuclei were pelleted via centrifugation at 2000 rpm and 4°C for 5 minutes, then resuspended in 600 µL cold nuclear extraction buffer. Biomag Plus Concanavalin A beads (Bangs Laboratories) were activated via two washes with 1 mL cold binding buffer (20 mM HEPES-KOH pH 7.5, 10 mM KCl, 1 mM CaCl_2_, & 1 mM MnCl_2_) using a magnetic rack, with 150 µL bead slurry used for each sample of 0.5 x 10^6^ cells. After the final wash, beads were resuspended in 300 µL cold binding buffer. While gently vortexing, nuclei were added to activated beads and then incubated on a rotating platform at 4°C for 10 minutes to facilitate nuclei binding. Nuclei:bead samples were resuspended in 1 mL blocking buffer (20 mM HEPES pH 7.5, 150 mM NaCl, 0.5 mM spermidine, 0.1% BSA, 2 mM EDTA, & protease inhibitors) and incubated at room temperature for 5 minutes. After blocking, samples were washed with 1 mL wash buffer (20 mM HEPES pH 7.5, 150 mM NaCl, 0.5 mM spermidine, 0.1% BSA, & protease inhibitors) then resuspended in 250 µL wash buffer. While gently vortexing, 5 µg of anti-H3K27me3 (#9733S, Cell Signaling Technology) in 250 µL wash buffer was added to each sample. Antibody binding was facilitated by incubating samples on a rotating platform at 4°C overnight. Samples were washed twice with 1 mL cold wash buffer and then resuspended in 250 µL cold wash buffer. While gently vortexing, 2.5 µL of CUTANA pAG-MNase (EpiCypher) was added, and samples were incubated at 4°C for 1 hour to facilitate binding. After incubation, samples were washed twice with 1 mL cold wash buffer and resuspended in 50 µL cold wash buffer. While gently vortexing, 3 µL 0.1 M CaCl_2_ was added to each sample. Samples were briefly flicked to mix and then incubated on ice for 30 minutes after which the pAG-MNase was inactivated by adding 50 µL 2x STOP buffer (340 mM NaCl, 20 mM EDTA, 4 mM EGTA, & 0.05% digitonin) supplemented with RNase A (ThermoFisher Scientific), proteinase K (ThermoFisher Scientific), glycogen (VWR), and spike-in DNA generated from MNase digestion of *Drosophila* S2 cells. Released fragments were cleaned up by incubating samples at 37°C for 20 minutes, followed by centrifugation at 16000xg and 4°C for 5 minutes. The resulting supernatant was mixed with 3 µL 10% SDS and 2.5 µL proteinase K solution, followed by incubation at 70°C for 10 minutes. To purify MNase-digested DNA fragments, samples underwent phenol-chloroform extraction. The final resulting aqueous phase was transferred and incubated with 2 µg glycogen and 750 µL ethanol at −20°C overnight. DNA was pelleted by centrifugation at 16000xg, air-dried, and then resuspended in 0.1x TE buffer. DNA fragments were checked for quality using the Qubit dsDNA HS Assay kit (Invitrogen) and TapeStation High Sensitivity DNA assay kit (Agilent).

Sequencing libraries of DNA fragments isolated following CUT&RUN were prepared using the NEBNext Ultra II DNA Library Prep Kit for Illumina (New England BioLabs) and the NEBNext Multiplex Oligos for Illumina (96 Unique Dual Index Primer Pairs; New England BioLabs). Libraries were checked for quality using the Qubit dsDNA HS Assay kit and TapeStation High Sensitivity DNA assay kit. Sequencing was performed using a NovaSeq 6000 with Illumina 150 bp paired-end reads at the Genomics Shared Resource, University of Colorado Cancer Center.

### Chromatin Immunoprecipitation (ChIP)

Native ChIP was performed with 2×10^6^ cells as previously described [37]. Cells were trypsinized, washed and subjected to hypotonic lysis (50 mM TrisHCl pH 7.4, 1 mM CaCl_2_, 0.2% Triton X-100, 10 mM NaButyrate, and protease inhibitor cocktail (Roche)) with micrococcal nuclease for 5 min at 37 °C to recover mono-to tri-nucleosomes. Nuclei were lysed by brief sonication and dialyzed into RIPA buffer (10 mM Tris pH 7.6, 1 mM EDTA, 0.1% SDS, 0.1% Na-Deoxycholate, 1% Triton X-100) for 2 hr at 4 °C. Soluble material was incubated with 3 μg of antibody bound to 50 μl protein A Dynabeads (Invitrogen) and incubated overnight at 4°C, with 5% reserved as input DNA. Magnetic beads were washed as follows: 3x RIPA buffer, 2x RIPA buffer + 300 mM NaCl, 2x LiCl buffer (250 mM LiCl, 0.5% NP-40, 0.5% NaDeoxycholate), 1x TE + 50 mM NaCl. Chromatin was eluted and treated with RNaseA and Proteinase K. ChIP DNA was purified and dissolved in H_2_O.

### ChIP-seq Library Preparation

ChIP-seq libraries were prepared from 5-10 ng ChIP DNA following the Illumina TruSeq protocol. The quality of the libraries was assessed using a D1000 ScreenTape on a 2200 TapeStation (Agilent) and quantified using a Qubit dsDNA HS Assay Kit (Thermo Fisher). Libraries with unique adaptor barcodes were multiplexed and sequenced on an Illumina NextSeq 500 (paired-end, 33 base pair reads). Typical sequencing depth was at least 20 million reads per sample.

### CUT&Tag

Genome-wide H3K27me3 domain changes were profiled using the CUT&Tag protocol adapted from EpiCypher. Cells were harvested via 0.25% trypsin-EDTA solution and counted using 0.4% trypan blue stain and Countess automated cell counter. A single-cell suspension was prepared by washing cells with cold 1x PBS and resuspending 1.0 x 10^6^ cells in 1 mL cold nuclear extraction buffer (20 mM HEPES-KOH pH 7.9, 10 mM KCl, 0.1% Triton x-100, 20% glycerol, & 0.5 mM spermidine), then incubated on ice for 10 minutes to isolate nuclei. Biomag Plus Concanavalin A beads were activated with two washes of cold bead activation buffer (20 mM HEPES-KOH pH 7.9, 10 mM KCl, 1 mM CaCl_2_, & 1 mM MnCl2_2_). After activation, beads were added to aliquots of 0.1 x 10^6^ isolated nuclei and incubated at room temperature for 10 minutes. Nuclei:bead samples were resuspended in 50 µL cold Antibody150 buffer (20 mM HEPES-KOH pH 7.5, 150 mM NaCl, 0.5 mM spermidine, 0.01% digitonin, 2 mM EDTA, & protease inhibitors) followed by the addition of 0.5 µg of anti-H3K27me3 (#9733S, Cell Signaling Technology). To facilitate primary antibody binding, samples were incubated on nutator at room temperature for 1 hour. Samples were resuspended in 50 µL Digitonin150 buffer (20 mM HEPES-KOH pH 7.5, 150 mM NaCl, 0.5 mM spermidine, 0.01% digitonin, & protease inhibitors) followed by the addition of 0.5 µg CUTANA anti-rabbit secondary antibody (#13-1047, EpiCypher). Samples were then incubated on nutator at room temperature for 1 hour to facilitate binding, followed by two washes with 200 µL cold Digitonin300 buffer (20 mM HEPES-KOH pH 7.5, 300 mM NaCl, 0.5 mM spermidine, 0.01% digitonin, & protease inhibitors). The tagmentation reaction was performed by resuspending samples in 50 µL cold Tagmentation buffer (20 mM HEPES pH 7.5, 300 mM NaCl, 0.5 mM spermidine, 10 mM MgCl_2_, & protease inhibitors) and incubating at 37°C in thermocycler for 1 hour. Samples were washed with 50 µL TAPS buffer (10 mM TAPS pH 8.5 & 0.2 mM EDTA) then resuspended in 5 µL SDS Release buffer (10 mM TAPS pH 8.5 & 0.1% SDS) to quench the tagmentation reaction and release adaptor-tagged DNA fragments into solution. After incubating samples at 58°C in thermocycler for 1 hour, 15 µL SDS Quench buffer (0.67% Triton x-100), 2 µL i5 universal primer (EpiCypher), 2 µL i7 barcoded primer (EpiCypher), 0.005 ng *Escherichia coli* spike-in DNA (made in-house), and 25 µL NEBNExt High Fidelity 2x PCR Master Mix were added to each sample for library construction with the following PCR protocol: 5 minutes at 58°C, 5 minutes at 72°C, 45 seconds at 98°C, 14 cycles of 15 seconds at 98°C and 10 seconds at 60°C, with a final extension for 1 minute at 72°C.

Libraries were checked for quality using the Qubit dsDNA HS Assay kit and TapeStation High Sensitivity DNA assay kit and then sequencing was performed using Novaseq 6000 (Novogene) to generate 150 bp paired-end reads.

### CUT&Tag in SU-DIPG-IV cells

Nuclei were prepared from SU-DIPG-IV cells, and up to 5×10^5^ cells were used per individual CUT&Tag reaction using CUT&Tag V3 (https://www.protocols.io/view/cut-amp-tag-direct-for-whole-cells-with-cutac-x54v9mkmzg3e/v3). pAG-Tn5 was purchased from Epicypher (Durham, North Carolina), and H3K27me3 antibody was purchased from Cell Signalling Technologies (#9733). Rabbit Isotype IgG control (ab172730) was purchased from Abcam. Antibodies were used at 1:100 dilution. Sequencing was performed on an Illumina NextSeq 500 using a P1/100 cycle kit with 50bp paired-end reads, targeting 10 million reads per sample.

### Data Analysis

CUT&Tag and CUT&RUN sequencing reads were trimmed using Cutadapt [38]: Illumina adapter sequences were removed and reads trimmed to 140 bp. Reads less than 35 bp were discarded. Trimmed reads were aligned to either the mm10 version of *Mus musculus* genome or the hg38 version of the human genome using bowtie2. Samtools [39] and bedtools [40] were used for processing aligned reads from sam to bed files. Duplicate reads were discarded for further analysis if the reads had the same start and end coordinates. Coverage at 100 bp windows genome-wide was calculated as the number of reads that mapped at that window, normalized by the factor N:

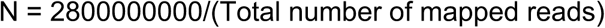

2800000000 was a number chosen arbitrarily in the range of total mapped base pairs in M. musculus/human genome. The normalized read counts were then smoothed with a running average spanning +/- 1000 bp around each 100 bp bin. The distribution of normalized read counts in 100 bp windows genome-wide was generated, and a “domain cutoff” was determined as the normalized read count that is greater than the normalized read count of F% of the windows. F was set based on background signal as 95 for all datasets except for SU-DIPG-IV datasets, where it was set as 75. Domains were called by linking adjacent windows with a normalized read count ≥ domain cutoff. To account for short disruptions due to issues of mappability, jumps of up to 750 bp were allowed while linking windows. A log2 ratio of H3K27me3 enrichment over IgG enrichment was calculated for all the putative domains. Those domains with a log2 enrichment greater than 2 (4-fold enrichment over IgG) were used for all downstream analyses. For performing disjoin and reduce operations, the GenomicRanges package in R was used [41].

For defining H3K27me3 domains from ENCODE, the following SEGWAY bed files were used. Control group: ENCFF489OJS.bigBed (A549), ENCFF248VPB.bigBed (HepG2), ENCFF097QCC.bigBed (K562), ENCFF412RIG.bigBed (SK-N-SH), ENCFF311LMS.bigBed (Karpas-422), ENCFF078XFK.bigBed (MM.1S), and ENCFF437JRM.bigBed (PC-3); Rb group: ENCFF835IDW.bigBed (WERI-Rb-1) and ENCFF137OLX.bigBed (HeLa-S3); CDKN2A group: ENCFF512MOL.bigBed (A673), ENCFF253DDW.bigBed (Panc1), ENCFF427CTV.bigBed (SJSA1), and ENCFF830MUH.bigBed (MCF-7).

## References

1. Li M, Liu GH, Izpisua Belmonte JC. Navigating the epigenetic landscape of pluripotent stem cells. Nat Rev Mol Cell Biol. 2012;13(8):524–35. Epub 2012/07/24. doi: 10.1038/nrm3393. PubMed PMID: 22820889.

2. Pengelly AR, Copur O, Jackle H, Herzig A, Muller J. A histone mutant reproduces the phenotype caused by loss of histone-modifying factor Polycomb. Science. 2013;339(6120):698-9. Epub 2013/02/09. doi: 10.1126/science.1231382. PubMed PMID: 23393264.

3. Kim JJ, Kingston RE. Context-specific Polycomb mechanisms in development. Nat Rev Genet. 2022;23(11):680–95. Epub 2022/06/11. doi: 10.1038/s41576-022-00499-0. PubMed PMID: 35681061; PubMed Central PMCID: PMCPMC9933872.

4. Bowman SK, Deaton AM, Domingues H, Wang PI, Sadreyev RI, Kingston RE, et al. H3K27 modifications define segmental regulatory domains in the Drosophila bithorax complex. Elife. 2014;3:e02833. Epub 2014/08/02. doi: 10.7554/eLife.02833. PubMed PMID: 25082344; PubMed Central PMCID: PMCPMC4139060.

5. Atlasi Y, Stunnenberg HG. The interplay of epigenetic marks during stem cell differentiation and development. Nat Rev Genet. 2017;18(11):643–58. Epub 2017/08/15. doi: 10.1038/nrg.2017.57. PubMed PMID: 28804139.

6. Alabert C, Barth TK, Reveron-Gomez N, Sidoli S, Schmidt A, Jensen ON, et al. Two distinct modes for propagation of histone PTMs across the cell cycle. Genes Dev. 2015;29(6):585–90. Epub 2015/03/21. doi: 10.1101/gad.256354.114. PubMed PMID: 25792596; PubMed Central PMCID: PMCPMC4378191.

7. Farrell JA, O’Farrell PH. From egg to gastrula: how the cell cycle is remodeled during the Drosophila mid-blastula transition. Annu Rev Genet. 2014;48:269–94. Epub 2014/09/10. doi: 10.1146/annurev-genet-111212-133531. PubMed PMID: 25195504; PubMed Central PMCID: PMCPMC4484755.

8. Kolodziejczyk AA, Kim JK, Tsang JC, Ilicic T, Henriksson J, Natarajan KN, et al. Single Cell RNA-Sequencing of Pluripotent States Unlocks Modular Transcriptional Variation. Cell Stem Cell. 2015;17(4):471–85. Epub 2015/10/03. doi: 10.1016/j.stem.2015.09.011. PubMed PMID: 26431182; PubMed Central PMCID: PMCPMC4595712.

9. Ter Huurne M, Chappell J, Dalton S, Stunnenberg HG. Distinct Cell-Cycle Control in Two Different States of Mouse Pluripotency. Cell Stem Cell. 2017;21(4):449–55 e4. doi: 10.1016/j.stem.2017.09.004. PubMed PMID: 28985526; PubMed Central PMCID: PMCPMC5658514.

10. Schlesinger S, Meshorer E. Open Chromatin, Epigenetic Plasticity, and Nuclear Organization in Pluripotency. Dev Cell. 2019;48(2):135–50. Epub 2019/01/30. doi: 10.1016/j.devcel.2019.01.003. PubMed PMID: 30695696.

11. Ying QL, Wray J, Nichols J, Batlle-Morera L, Doble B, Woodgett J, et al. The ground state of embryonic stem cell self-renewal. Nature. 2008;453(7194):519-23. Epub 2008/05/24. doi: 10.1038/nature06968. PubMed PMID: 18497825; PubMed Central PMCID: PMCPMC5328678.

12. Kuroda MI, Kang H, De S, Kassis JA. Dynamic Competition of Polycomb and Trithorax in Transcriptional Programming. Annu Rev Biochem. 2020;89:235–53. Epub 2020/01/14. doi: 10.1146/annurev-biochem-120219-103641. PubMed PMID: 31928411; PubMed Central PMCID: PMCPMC7311296.

13. Streubel G, Watson A, Jammula SG, Scelfo A, Fitzpatrick DJ, Oliviero G, et al. The H3K36me2 Methyltransferase Nsd1 Demarcates PRC2-Mediated H3K27me2 and H3K27me3 Domains in Embryonic Stem Cells. Mol Cell. 2018;70(2):371–9 e5. Epub 2018/04/03. doi: 10.1016/j.molcel.2018.02.027. PubMed PMID: 29606589.

14. Fang Y, Tang Y, Zhang Y, Pan Y, Jia J, Sun Z, et al. The H3K36me2 methyltransferase NSD1 modulates H3K27ac at active enhancers to safeguard gene expression. Nucleic Acids Res. 2021;49(11):6281–95. Epub 2021/06/10. doi: 10.1093/nar/gkab473. PubMed PMID: 34107030; PubMed Central PMCID: PMCPMC8216457.

15. Chen H, Hu B, Horth C, Bareke E, Rosenbaum P, Kwon SY, et al. H3K36 dimethylation shapes the epigenetic interaction landscape by directing repressive chromatin modifications in embryonic stem cells. Genome Res. 2022;32(5):825–37. Epub 2022/04/10. doi: 10.1101/gr.276383.121. PubMed PMID: 35396277; PubMed Central PMCID: PMCPMC9104706.

16. Kumar B, Elsasser SJ. Quantitative Multiplexed ChIP Reveals Global Alterations that Shape Promoter Bivalency in Ground State Embryonic Stem Cells. Cell Rep. 2019;28(12):3274–84 e5. Epub 2019/09/19. doi: 10.1016/j.celrep.2019.08.046. PubMed PMID: 31533047; PubMed Central PMCID: PMCPMC6859498.

17. Tse AN, Rendahl KG, Sheikh T, Cheema H, Aardalen K, Embry M, et al. CHIR-124, a novel potent inhibitor of Chk1, potentiates the cytotoxicity of topoisomerase I poisons in vitro and in vivo. Clin Cancer Res. 2007;13(2 Pt 1):591-602. Epub 2007/01/27. doi: 10.1158/1078-0432.CCR-06-1424. PubMed PMID: 17255282.

18. Classon M, Harlow E. The retinoblastoma tumour suppressor in development and cancer. Nat Rev Cancer. 2002;2(12):910–7. Epub 2002/12/03. doi: 10.1038/nrc950. PubMed PMID: 12459729.

19. Foulkes WD, Flanders TY, Pollock PM, Hayward NK. The CDKN2A (p16) gene and human cancer. Mol Med. 1997;3(1):5–20. Epub 1997/01/01. PubMed PMID: 9132280; PubMed Central PMCID: PMCPMC2230107.

20. Hoffman MM, Buske OJ, Wang J, Weng Z, Bilmes JA, Noble WS. Unsupervised pattern discovery in human chromatin structure through genomic segmentation. Nat Methods. 2012;9(5):473–6. Epub 2012/03/20. doi: 10.1038/nmeth.1937. PubMed PMID: 22426492; PubMed Central PMCID: PMCPMC3340533.

21. Luo Y, Hitz BC, Gabdank I, Hilton JA, Kagda MS, Lam B, et al. New developments on the Encyclopedia of DNA Elements (ENCODE) data portal. Nucleic Acids Res. 2020;48(D1):D882-D9. Epub 2019/11/13. doi: 10.1093/nar/gkz1062. PubMed PMID: 31713622; PubMed Central PMCID: PMCPMC7061942.

22. Liu I, Jiang L, Samuelsson ER, Marco Salas S, Beck A, Hack OA, et al. The landscape of tumor cell states and spatial organization in H3-K27M mutant diffuse midline glioma across age and location. Nat Genet. 2022;54(12):1881–94. Epub 2022/12/06. doi: 10.1038/s41588-022-01236-3. PubMed PMID: 36471067; PubMed Central PMCID: PMCPMC9729116 Medicines. M.N. is Scientific Advisor for 10X Genomics. M.M. is a SAB member for Cygnal Therapeutics. M.L. Suva is an equity holder, scientific cofounder and advisory board member of Immunitas Therapeutics. K.L.L. is the founder and equity holder of Travera and receives consulting fees from BMS, Integragen, Rarecyte and research support from Lilly, BMS and Amgen. J.S. is now (but not when contributing to this manuscript) an employee of 10X Genomics. The remaining authors declare no competing interests.

23. Harutyunyan AS, Krug B, Chen H, Papillon-Cavanagh S, Zeinieh M, De Jay N, et al. H3K27M induces defective chromatin spread of PRC2-mediated repressive H3K27me2/me3 and is essential for glioma tumorigenesis. Nat Commun. 2019;10(1):1262. Epub 2019/03/21. doi: 10.1038/s41467-019-09140-x. PubMed PMID: 30890717; PubMed Central PMCID: PMCPMC6425035.

24. Toogood PL, Harvey PJ, Repine JT, Sheehan DJ, VanderWel SN, Zhou H, et al. Discovery of a potent and selective inhibitor of cyclin-dependent kinase 4/6. J Med Chem. 2005;48(7):2388–406. Epub 2005/04/02. doi: 10.1021/jm049354h. PubMed PMID: 15801831.

25. Zerihun MB, Vaillant C, Jost D. Effect of replication on epigenetic memory and consequences on gene transcription. Phys Biol. 2015;12(2):026007. Epub 2015/04/18. doi: 10.1088/1478-3975/12/2/026007. PubMed PMID: 25884278.

26. Seller CA, Cho CY, O’Farrell PH. Rapid embryonic cell cycles defer the establishment of heterochromatin by Eggless/SetDB1 in Drosophila. Genes Dev. 2019;33(7-8):403–17. Epub 2019/02/28. doi: 10.1101/gad.321646.118. PubMed PMID: 30808658; PubMed Central PMCID: PMCPMC6446540.

27. Yoon H, Kim TS, Braciale TJ. The cell cycle time of CD8+ T cells responding in vivo is controlled by the type of antigenic stimulus. PLoS One. 2010;5(11):e15423. Epub 20101108. doi: 10.1371/journal.pone.0015423. PubMed PMID: 21079741; PubMed Central PMCID: PMCPMC2975678.

28. Beguelin W, Popovic R, Teater M, Jiang Y, Bunting KL, Rosen M, et al. EZH2 is required for germinal center formation and somatic EZH2 mutations promote lymphoid transformation. Cancer Cell. 2013;23(5):677–92. Epub 2013/05/18. doi: 10.1016/j.ccr.2013.04.011. PubMed PMID: 23680150; PubMed Central PMCID: PMCPMC3681809.

29. Jain SU, Rashoff AQ, Krabbenhoft SD, Hoelper D, Do TJ, Gibson TJ, et al. H3 K27M and EZHIP Impede H3K27-Methylation Spreading by Inhibiting Allosterically Stimulated PRC2. Mol Cell. 2020;80(4):726–35 e7. Epub 20201012. doi: 10.1016/j.molcel.2020.09.028. PubMed PMID: 33049227; PubMed Central PMCID: PMCPMC7680438.

30. Sarthy JF, Meers MP, Janssens DH, Henikoff JG, Feldman H, Paddison PJ, et al. Histone deposition pathways determine the chromatin landscapes of H3.1 and H3.3 K27M oncohistones. Elife. 2020;9. Epub 2020/09/10. doi: 10.7554/eLife.61090. PubMed PMID: 32902381; PubMed Central PMCID: PMCPMC7518889.

31. Krishan A. Rapid flow cytofluorometric analysis of mammalian cell cycle by propidium iodide staining. J Cell Biol. 1975;66(1):188–93. Epub 1975/07/01. doi: 10.1083/jcb.66.1.188. PubMed PMID: 49354; PubMed Central PMCID: PMCPMC2109516.

32. Shechter D, Dormann HL, Allis CD, Hake SB. Extraction, purification and analysis of histones. Nat Protoc. 2007;2(6):1445–57. Epub 2007/06/05. doi: 10.1038/nprot.2007.202. PubMed PMID: 17545981.

33. Schneider CA, Rasband WS, Eliceiri KW. NIH Image to ImageJ: 25 years of image analysis. Nat Methods. 2012;9(7):671–5. Epub 2012/08/30. doi: 10.1038/nmeth.2089. PubMed PMID: 22930834; PubMed Central PMCID: PMCPMC5554542.

34. Sidoli S, Kori Y, Lopes M, Yuan ZF, Kim HJ, Kulej K, et al. One minute analysis of 200 histone posttranslational modifications by direct injection mass spectrometry. Genome Res. 2019;29(6):978–87. Epub 2019/05/28. doi: 10.1101/gr.247353.118. PubMed PMID: 31123082; PubMed Central PMCID: PMCPMC6581051.

35. Karch KR, Sidoli S, Garcia BA. Identification and Quantification of Histone PTMs Using High-Resolution Mass Spectrometry. Methods Enzymol. 2016;574:3–29. Epub 2016/07/18. doi: 10.1016/bs.mie.2015.12.007. PubMed PMID: 27423855; PubMed Central PMCID: PMCPMC5089704.

36. Hainer SJ, Fazzio TG. High-Resolution Chromatin Profiling Using CUT&RUN. Curr Protoc Mol Biol. 2019;126(1):e85. Epub 2019/01/29. doi: 10.1002/cpmb.85. PubMed PMID: 30688406; PubMed Central PMCID: PMCPMC6422702.

37. Martire S, Gogate AA, Whitmill A, Tafessu A, Nguyen J, Teng YC, et al. Phosphorylation of histone H3.3 at serine 31 promotes p300 activity and enhancer acetylation. Nat Genet. 2019;51(6):941–6. Epub 2019/06/04. doi: 10.1038/s41588-019-0428-5. PubMed PMID: 31152160; PubMed Central PMCID: PMCPMC6598431.

38. Martin M. Cutadapt removes adapter sequences from high-throughput sequencing reads. 2011. 2011;17(1):3. Epub 2011-08-02. doi: 10.14806/ej.17.1.200.

39. Li H, Handsaker B, Wysoker A, Fennell T, Ruan J, Homer N, et al. The Sequence Alignment/Map format and SAMtools. Bioinformatics. 2009;25(16):2078–9. Epub 2009/06/10. doi: 10.1093/bioinformatics/btp352. PubMed PMID: 19505943; PubMed Central PMCID: PMCPMC2723002.

40. Quinlan AR, Hall IM. BEDTools: a flexible suite of utilities for comparing genomic features. Bioinformatics. 2010;26(6):841–2. Epub 2010/01/30. doi: 10.1093/bioinformatics/btq033. PubMed PMID: 20110278; PubMed Central PMCID: PMCPMC2832824.

41. Lawrence M, Huber W, Pages H, Aboyoun P, Carlson M, Gentleman R, et al. Software for computing and annotating genomic ranges. PLoS Comput Biol. 2013;9(8):e1003118. Epub 2013/08/21. doi: 10.1371/journal.pcbi.1003118. PubMed PMID: 23950696; PubMed Central PMCID: PMCPMC3738458.

